# circRNA-sponging: a pipeline for extensive analysis of circRNA expression and their role in miRNA sponging

**DOI:** 10.1101/2023.01.19.524495

**Authors:** Markus Hoffmann, Leon Schwartz, Octavia-Andreea Ciora, Nico Trummer, Lina-Liv Willruth, Jakub Jankowski, Hye Kyung Lee, Jan Baumbach, Priscilla Furth, Lothar Hennighausen, Markus List

## Abstract

**Motivation:** Circular RNAs (circRNAs) are long non-coding RNAs (lncRNAs) often associated with diseases and considered potential biomarkers for diagnosis and treatment. Among other functions, circRNAs have been shown to act as microRNA (miRNA) sponges, preventing the role of miRNAs that repress their targets. However, there is no pipeline to systematically assess the sponging potential of circRNAs.

**Results:** We developed circRNA-sponging, a nextflow pipeline that (1) identifies circRNAs via backsplicing junctions detected in RNA-seq data, (2) quantifies their expression values in relation to their linear counterparts spliced from the same gene, (3) performs differential expression analysis, (4) identifies and quantifies miRNA expression from miRNA-sequencing (miRNA-seq) data, (5) predicts miRNA binding sites on circRNAs, (6) systematically investigates potential circRNA-miRNA sponging events, (7) creates a network of competing endogenous RNAs, and (8) identifies potential circRNA biomarkers. We showed the functionality of the circRNA-sponging pipeline using RNA sequencing data from brain tissues, where we identified two distinct types of circRNAs characterized by a specific ratio of the number of the binding site to the length of the transcript. The circRNA-sponging pipeline is the first end-to-end pipeline to identify circRNAs and their sponging systematically with raw total RNA-seq and miRNA-seq files, allowing us to better indicate the functional impact of circRNAs as a routine aspect in transcriptomic research.

**Availability:** https://github.com/biomedbigdata/circRNA-sponging

**Supplementary Material:** Supplementary data are available at Bioinformatic Advances online.

## INTRODUCTION

Circular RNAs (circRNAs) are classified as long non-coding RNAs (lncRNAs), even though a few have been reported to encode proteins [1]. circRNAs are characterized by their loop structure, which makes them less prone to degradation [2,3]. The biogenesis of circRNAs is explained by the occurrence of a backsplicing event (see Suppl. Fig. 1) during the alternative splicing process of precursor messenger RNA (pre-mRNA), where the 5’ terminus of an upstream exon and the 3’ terminus of a downstream exon are covalently joined [2]. The difference between circRNAs and linear RNAs is the lack of a 5’ cap and a 3’ polyadenylation (poly(A)) tail along with its circular shape, which makes circRNAs more stable and, in most cases, resistant to exonuclease activity [4–6]. These circular molecules can be made up of exonic and intronic regions of its spliced pre-mRNA and are thus found in a variety of sizes, ranging from 100 to >4,000 nucleotides [4,7]. circRNAs are conserved across species, and their expression is tissue- and disease-specific [3,8,9]. Hence, they can play an important role as biomarkers and therapeutic targets [9–11]. Another type of non-coding RNA is microRNA (miRNA) which plays a role in post-transcriptional gene regulation [12,13] and is involved in many biological processes and diseases [14]. miRNAs bind their target genes via the RNA-induced silencing complex causing their degradation or preventing their translation [15].

The possible interplay between circRNAs, miRNAs, messenger RNAs (mRNAs) that code for proteins, and other types of RNA that share miRNA binding sites gives rise to a large regulatory network. Salmena et al. proposed that any RNA that carries miRNA binding sites (e.g., mRNAs, circRNAs, pseudogenes, transcripts of 3’ untranslated regions (UTRs), and lncRNAs) can act as a competing endogenous RNA (ceRNA) [16] that competes for the limited pool of available miRNAs in a cell. As a result of this competition, an overexpressed RNA can sponge away miRNAs required for the regulation of other RNAs, which can explain why non-coding RNAs, such as circRNAs, can be implicated in a phenotype.

The enhanced stability of circRNAs might allow them to work as buffers for miRNAs by binding them until sufficient miRNAs are present to outnumber the circRNA binding sites [9]. The regulatory function of circRNAs and their alleged association with diseases are the main reasons why identifying sponging activity between circRNAs and miRNAs is of particular interest. The presence of an interaction between miRNAs and circRNAs has been repeatedly proven, and several circRNAs (e.g., CDR1as/CiRS-7, SRY [17], and circNCX1 [18]) have been recognized as miRNA sponges. Even though individual studies confirmed the existence of circRNA sponges, further studies are needed to elucidate the role of circRNAs in miRNA-mediated gene regulation.

From a computational point of view, the detection of circRNAs is difficult due to their circular shape and the lack of poly(A) tail, which makes it unlikely to observe them in poly(A)-enriched RNA sequencing (RNA-seq) libraries [7]. Hence, circRNAs can only be robustly detected in libraries without poly(A) enrichment, such as ribosomal RNA (rRNA) depleted RNA-seq and total RNA-sequencing (total RNA-seq), which do not deplete circRNAs [7]. Identification of circRNAs relies on the detection of backsplicing junctions among the unmapped reads, which allows for the estimation of circRNA abundance. By focusing on the backsplicing junction alone, the expression of circRNAs in relation to their linear counterparts is typically underestimated [19].

Several approaches for circRNA analysis have been proposed. Chen et al. reviewed 100 existing circRNA-related tools for circRNA detection, annotation, downstream analysis, as well as network analysis [20]. They list a total of 44 circRNA identification tools including, but not limited to, CIRCexplorer [21,22], find_circ [23], CIRI [24], KNIFE [25], and circRNA_finder [26]. They also present a total of 14 circRNA annotation databases collecting circRNA information from the literature, such as circBase [27] and CIRCpedia [21,28]. Other circRNA-related tools include databases for feature collection and storing circRNA information related to disease and biomarkers. In addition, circRNA network identification tools model the interactions between circRNAs and miRNAs, lncRNAs, or RNA-binding proteins. Other tools for downstream analysis of circRNAs cover alternative splicing detection, circRNA assembly, and structure prediction and visualization [20]. To the best of our knowledge, none of the tools provides a comprehensive and automated circRNA sponging analysis integrating identification and quantification of both circRNAs and miRNAs, a systematic investigation of potential circRNA-miRNA sponging events, and a ceRNA network analysis. We developed “circRNA-sponging”, a nextflow pipeline integrating state-of-the-art methods to (1) detect circRNAs via identifying backsplicing junctions from total RNA-seq data, (2) quantify their expression values relative to linear transcripts, (3) perform differential expression analysis, (4) identify and quantify miRNA expression from miRNA-sequencing (miRNA-seq) data, (5) predict miRNA binding sites on circRNAs, (6) systematically investigate potential circRNA-miRNA sponging events, (7) create a ceRNA network, and (8) identify potential circRNA biomarkers using the ceRNA network (Fig. 1).

**Figure 1:**
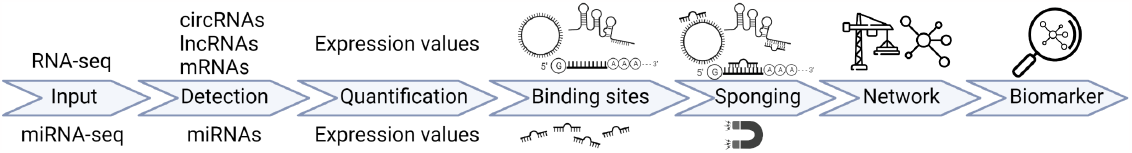
Overview of the individual steps of the pipeline. total RNA-seq data processing is shown on top, and miRNA-seq processing on the bottom. In the miRNA sponging step, these results are integrated for network analysis and biomarker detection.

We demonstrate the potential of the circRNA-sponging pipeline on matched rRNA-depleted RNA-seq and miRNA-seq data from mouse brain tissues.

## MATERIALS AND METHODS

### Data

Using circRNA-sponging, we processed a total of 23 samples of matched single-end rRNA-depleted RNA-seq and miRNA-seq data for four brain regions (cerebellum, cortex, hippocampus, olfactory bulb). Samples include 3 replicates for wild-type (WT) and 2-3 CDR1 knock-out (KO) mouse replicates (GEO accessions: GSE100265, GSE93129) [29]. (see Suppl. Table 1). We use the mm10 genome version for mapping.

### Pipeline Architecture

The circRNA-sponging pipeline is implemented in R (v. 4.2.0) and Python (v. 3.8.12) and wrapped with nextflow version 22.04.0.5697. The pipeline is hosted on dockerhub and will pull the required docker image when executed. The relevant image was built under docker version 22.06. It follows the nf-core guidelines [30] and encompasses several state-of-the-art techniques organized into three modules: (1) the circRNA module, (2) the miRNA module, and (3) the sponging module, the latter of which can only be performed if both other modules have been executed (Fig. 2). In the following, we provide a deeper insight into each module and highlight important components of the pipeline.

1. *The circRNA module* addresses the identification, quantification, and miRNA binding site prediction of circRNAs. For read mapping, we employ the STAR [31] aligner, which provides support for the detection of splice-junction and fusion reads. The resulting unmapped split-reads are used by CIRCexplorer2, which uses a combination of methods (i.e., a de novo assembly approach to identify novel circRNA and a reference-based approach, which uses known exon-exon junctions to map backsplicing events to known genes) to increase the accuracy of its predictions [21] to identify backsplicing events. We could confirm the excellent performance of CIRCexplorer2 using data simulated with polyester [32] from the linear mouse reference genome GRCm38 at varying sequencing depth, where we rarely detect false positive backsplicing junctions and zero false positive circRNAs (Suppl. Fig. 2a). Next, raw read counts are normalized with DESeq2 [33], and circRNAs with low expression levels are excluded to reduce false positives. By default, only circRNAs with a normalized read count >5 in at least 20% of samples are retained. Database annotation is performed using circBase [27], which covers curated circRNAs with experimental evidence of several model organisms. We use psirc [19] to quantify circRNA expression levels, as the detection of backsplicing junctions alone does not reflect circRNA expression levels in relation to the gene’s expression. To mitigate this, psirc employs kallisto [34] and considers both linear and circular transcripts in the expectation-maximization step to produce comparable expression values. psirc corrects for various sequencing biases that can affect circRNA detection, such as coverage bias, mapping bias, read length bias, and alternative splicing bias. Yu et al. showed that psirc provides a more accurate identification of circRNA expression levels by validating their method with experimental data [19]. If the data has been sampled from different conditions (e.g., case and control), the quantified linear transcripts and circRNAs can be used to perform a differential expression analysis using DESeq2 [33]. The pipeline generates heatmaps, volcano plots, and principal component analysis (PCA) of the circRNAs and linear transcripts between conditions. We analyze alternative splicing between circular and linear transcripts on a gene level using SUPPA2 [35]. In order to integrate circular transcripts, we construct a merged gene annotation file consisting of both linear and circular transcripts. Based on this input, we generate percent spliced-in (PSI) values for linear and circular isoforms with the SUPPA2 step psiPerIsoform [35]. We, additionally, normalized the linear and circular PSI values for a gene by their sample-wise mean to account for differences in overall linear and circular splicing frequencies. Both non-normalized and normalized PSI values are automatically visualized. To boost reliability, we predict circRNA-miRNA binding sites using a majority voting between miRanda [36], PITA [37], and TarPmiR [38] since each method has a distinct approach for predicting miRNA binding sites. Testing these tools with random miRNA sequences shows that up to 25% of the reported binding sites may be false positives (Suppl. Fig. 2b-d), which aligns with previous findings [39]. Thus, we consider a circRNA-miRNA binding site as relevant if it is predicted by at least two out of the three methods. miRanda considers seed matching, conservation, and free energy, and we consider predictions with a score above the 25% quantile. PITA additionally considers site accessibility and target-site abundance. TarPmiR further integrates machine learning to improve results for supported organisms [38]. We further incorporate experimentally validated target sites from DIANA-LncBase v3 [40], miRTarBase [41,42], and miRWalk3.0 [43].
2. *The miRNA module* covers the quantification and processing of miRNA expression. miRDeep2 [44] is used to obtain miRNA counts. Alternatively, already mapped miRNA expression data can be provided. Raw counts are normalized with DESeq2 [33] followed by a filtering step, where by default, miRNAs with a normalized read count > 5 in at least 20% of samples are retained.
3. *The sponging module* is used for the identification of crosstalk between circRNAs, miRNAs as well as ceRNA interactions of circRNAs with other transcripts. To identify potential sponging activity, we perform a correlation analysis of circRNA-miRNA pairs, where a negative correlation coefficient indicates a sponging relationship. For all circRNA-miRNA pairs (i.e., a circRNA that harbors at least one binding site for the miRNA), we compute a Pearson correlation coefficient along with the normalized residual sum of squares and the adjusted p-value after the Benjamini-Hochberg correction. Pairs are filtered (e.g., p-adjusted < 0.05, RMSE < 1.5, and optionally by the number of binding sites) and are considered potential sponging candidates. We further construct a ceRNA network using SPONGE [12] on matched gene and miRNA expression data. Finally, we apply spongEffects [45] to extract ceRNA modules consisting of circRNAs with a high node centrality score in the ceRNA network and their direct neighbors. For each module, spongEffects computes a sample-specific enrichment score (i.e., the spongEffects enrichment scores are calculated using one of three gene set enrichment approaches: Gene Set Enrichment Analysis (ssGSEA) and Gene Set Variation analysis (GSVA) algorithms as implemented in the GSVA package (version 1.34.0) [46], or Overall Expression (OE) [47]. These approaches can calculate spongEffects scores even if some genes in the ceRNAs modules are missing. The resulting module-by-sample score matrices can be used for further analysis. No major differences were observed between the three methods, and the choice of the optimal tool depends on the specific task and dataset [45], such as differential analysis between groups or supervised machine learning. The spongEffects scores are then used for training and testing a random model classifier to distinguish between groups of samples (e.g., healthy and control) [48]. The prediction power is then measured by a 5-fold cross-validation and a comparison to random modules (i.e., sampling modules of the same size as the ones predicted while preserving the size distribution of the real modules). In our example data set, we used 16 samples for training (four samples for cerebellum, cortex, hippocampus, and olfactory bulb) and eight samples for testing (two samples for cerebellum, cortex, hippocampus, and olfactory bulb).

**Figure 2:**
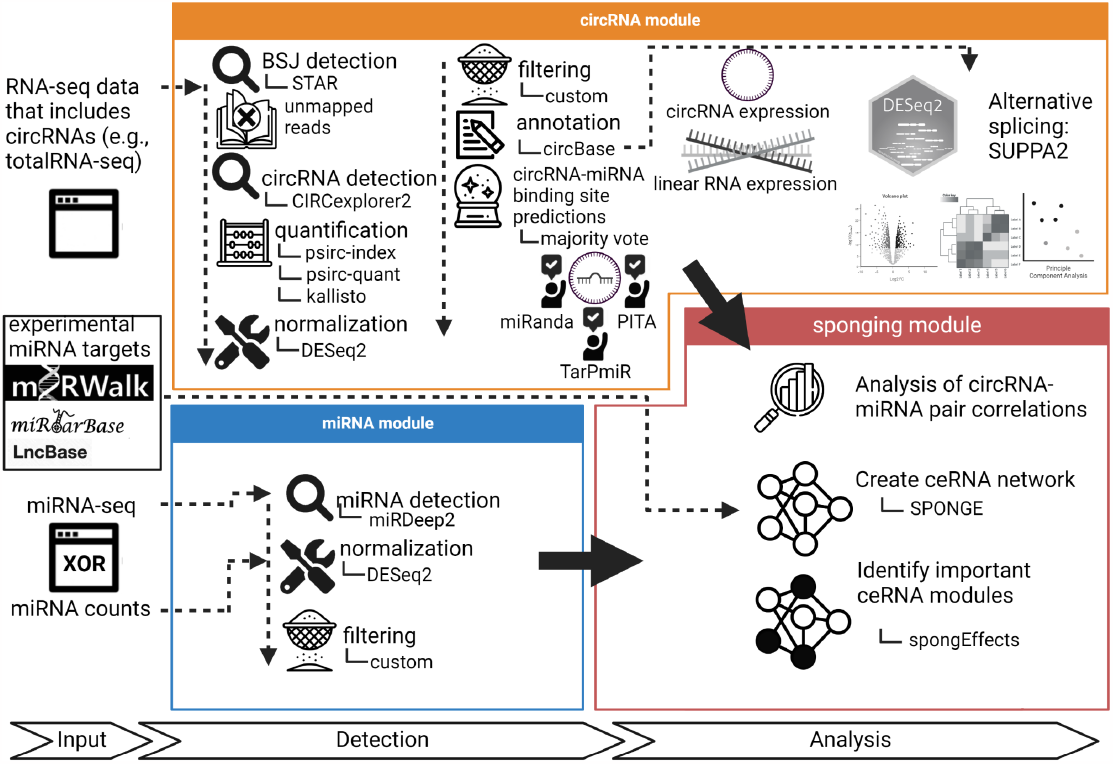
Workflow of the circRNA-sponging pipeline. The pipeline consists of three modules: (1) the circRNA module, (2) the miRNA module, and (3) the sponging module. In (1), we detect circRNAs via identifying backsplicing junctions from total RNA-seq data, quantify their expression values, perform a differential expression analysis, and predict miRNA binding sites on circRNAs using a majority vote between three state-of-the-art methods. In (2), we either detect and quantify miRNAs in raw miRNA-seq or directly process miRNA expression data. In (3), we systematically investigate circRNA-miRNA sponging events, create a ceRNA network and use it to identify potential circRNA biomarkers.

## RESULTS AND DISCUSSION

circRNAs are highly abundant and conserved in the mammalian brain [49,50]. To demonstrate the capabilities of the circRNA-sponging pipeline, we analyzed a public RNA-seq data set from the mouse brain. We focused on the sponging capacity of circRNAs and their potential role as ceRNAs.

### Comparing circRNA and host gene expression reveals changes in circRNA splicing

In total, we detected 46,380 and 27,390 circRNAs before and after filtering, respectively. This number aligns with the known high abundance of circRNAs in brain tissue [49,50]. We could annotate only 1,027 (∽4%) of the circRNAs that passed the filter (Suppl. Fig. 3a), as comparably few circRNAs have thus far been annotated in mice using circBase. psirc-estimated expression levels, which take reads mapping to parts other than the backsplicing junction of the circRNA into account, are up to 6-fold higher compared to counts derived from backsplicing junctions only (Fig. 3c, per tissue type: Suppl. Fig. 3d-g). We observed a generally higher expression of circRNAs in the cerebellum compared to other brain regions, which could indicate a higher importance of the circRNAs in this brain region (Suppl. Fig. 3b).

**Figure 3:**
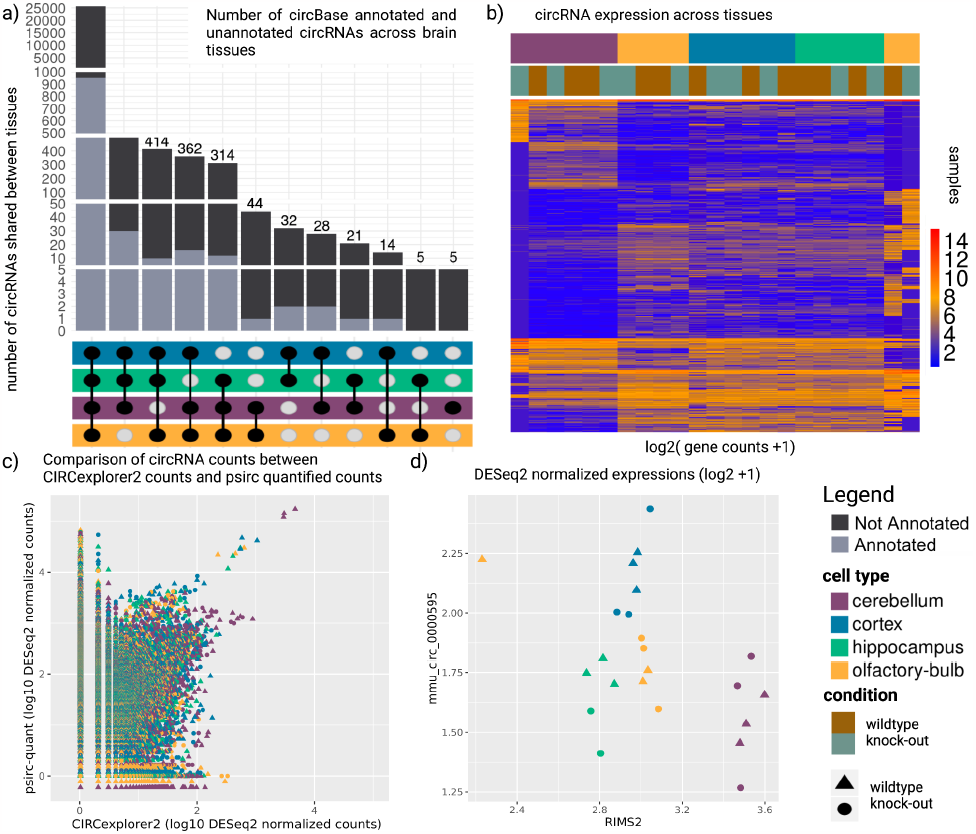
circRNA results of the mouse brain regions data set. a) circRNAs shared between brain regions. b) expression of circRNAs across tissues and experimental conditions. c) Comparison of psirc-quant quantified circRNA counts to CIRCexplorer2 counts. d) comparison between a circRNA originating from RIMS2 and expression of the linear RIMS2 gene.

Concerning the miRNA binding sites, we observed that with a higher number of binding sites, the Pearson correlation increases (Suppl. Fig. 4). Despite the high number of shared circRNAs across brain regions (Fig. 3a), we observed a brain region-specific abundance of overall circRNAs levels (Fig. 3b, Suppl. Fig. 3b). However, expression levels of the circRNAs differ considerably, such that samples cluster by brain region (Fig. 3b, Suppl. Fig. 3c). Rybak-Wolf et al. reported circRNAs of twelve host genes (TULP4, RIMS2, ELF2, PHF21A, MYST4, CDR1, STAU2, SV2B, CPSF6, DYM, RMST, and RTN4) [50]. They speculated on the importance of circRNAs originating from these genes for brain cell identity, but we posit that a change in circRNA expression alone does not necessarily imply a functional role as circRNA expression is coupled to the expression of the host gene, as we expect the number of reads mapping to the backsplicing junctions to correlate. We detected nine of the twelve circRNAs (all but TULP4, SV2B, and RMST) in our analysis (Suppl. Table 2, Suppl. Fig. 5a-b). The difference between KO and WT samples is negligible, with the exception of the CDR1 region (mmu_hsa_circ_0001878 in circBase annotation) that was targeted successfully (Suppl. Fig. 5c-d). When excluding the circRNA of the CDR1, we observed a clear separation between the cerebellum and other brain regions, while the cortex and hippocampus are more similar [51]. Our analysis revealed a total of 33 circRNAs that show significantly different expression between brain regions (p-adjusted < 0.01, absolute log2 fold change > 5, Suppl. Table 3). By comparing the expression level of the circRNAs to the linear transcripts, as facilitated by psirc-quant, we can identify cases where the expression of circRNAs increases beyond the level suggested by the overall gene expression. Such cases could offer evidence for the functional importance of a circRNA. For example, mmu_circ_0000595, a circRNA of host gene RIMS2, shows a higher expression in the cortex, whereas the expression of the host gene RIMS2 remains stable in this tissue (Fig. 3d, see also Suppl. Fig. 6a-p). We investigate the output of SUPPA2 to explore the relationship between the quantity of circular and linear transcripts per shared gene more thoroughly. As expected, the PSI values of the linear transcripts are mostly close to 100%, whereas circular RNAs are rare and show overall very low PSI values. However, there are instances where circular transcripts show high PSI values (Suppl. Fig. 7a). Differential splicing analysis of circular transcripts revealed that only very few are significantly differentially spliced between cell types, i.e., pass both filters of a change in PSI >= 25 % and a p-value of <= 0.01 (Suppl. Fig. 7b). We can observe that two circRNAs (chr15:34600014-34625031_- with its host genes HYDIN between cortex-hippocampus and chr8:110298074-110334816_+ with its host gene NIPAL2 between hippocampus-olfactory-bulb) are considered differentially spliced in (Suppl. Fig. 7). These results are similar for normalized PSI values (Suppl. Fig. 8), where we accounted for sample-specific differences. Our results thus suggest that the splicing ratio of linear to circular RNA expression does not change between different brain regions.

### A ceRNA network reveals circRNAs acting as miRNA sponges

If matched total RNA and miRNA sequencing data are provided, circRNA-sponging infers a ceRNA network using the R package SPONGE [12] and visualizes the result (Fig. 4). Important players in this regulatory network are characterized by a large node degree, i.e., they indirectly regulate many of the connected RNAs via sequestering miRNA copies. Since the network inferred by SPONGE does not offer insights into individual samples or conditions, we subsequently computed spongEffects [45] enrichment scores which capture the interaction of individual circRNAs and their target genes. As these scores are sample-specific, they can offer insights into condition-specific circRNA sponging activity. spongeEffects scores can also be used as features for machine learning tasks such as classification [45]. Since the number of available samples for training is rather small here, the random forest reported subset accuracy drops considerably on the holdout set in ten-fold cross-validation. While the cortex and hippocampus are difficult to differentiate, the cerebellum can be robustly distinguished from other brain regions (Suppl. Fig. 9). In particular, two circRNAs, *chr10:9770449-9800068_-* and *chr10:79860969-79862010_*+ stand out as distinctive features of the cerebellum (Suppl. Fig. 10-11). While the inferred ceRNA network shows that circRNAs in the mouse brain are regulatory active through miRNA sponging, a larger number of samples is likely needed to fully resolve brain region-specific sponging activity.

**Figure 4:**
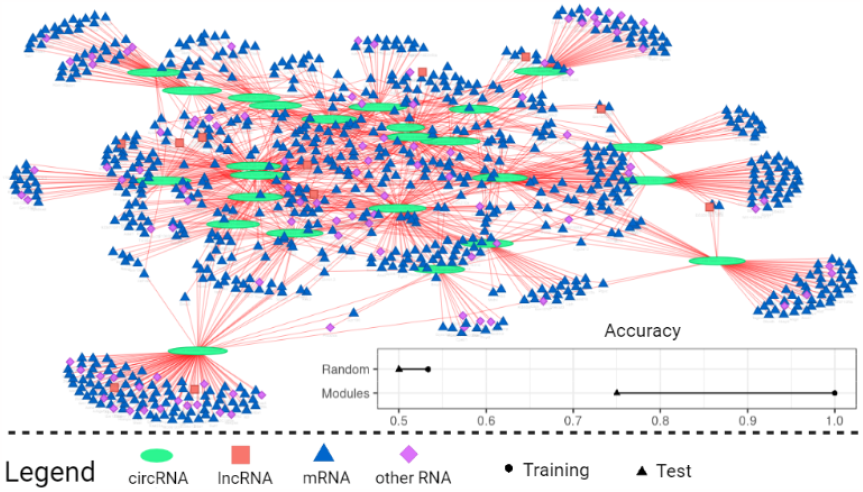
circRNA-ceRNA-subnetwork with the top 25 ceRNA modules ranked by the number of significant interactions (node degree in the network). For each ceRNA module consisting of the circRNA and its target genes, we computed spongEffects enrichment scores and used them as input for a random forest model. The bottom-right corner shows the subset accuracy of this model in distinguishing different brain regions on the training and test set. The results of a model trained on random modules of the same size show random performance.

### Comparing the number of circRNA binding sites with respect to their length reveals two distinct clusters

miRNA sponging has long been considered a potential function of circRNAs [17]. To fulfill this function, it would be beneficial for circRNAs to carry a large number of miRNA binding sites, and indeed, some known circRNAs, such as CDR1as harboring over 70 miRNA binding sites for miR-7 alone [52], fit the hypothesis well. To investigate if this is a general property of circRNAs, we systematically compared the length of a transcript to the number of binding sites, expecting to observe a larger ratio for circRNAs compared to the 3’ untranslated regions of linear transcripts, where miRNA binding sites are predominantly located (Fig. 5). While linear transcripts show a very diverse picture, circRNA length correlates well with the number of binding sites.

**Figure 5:**
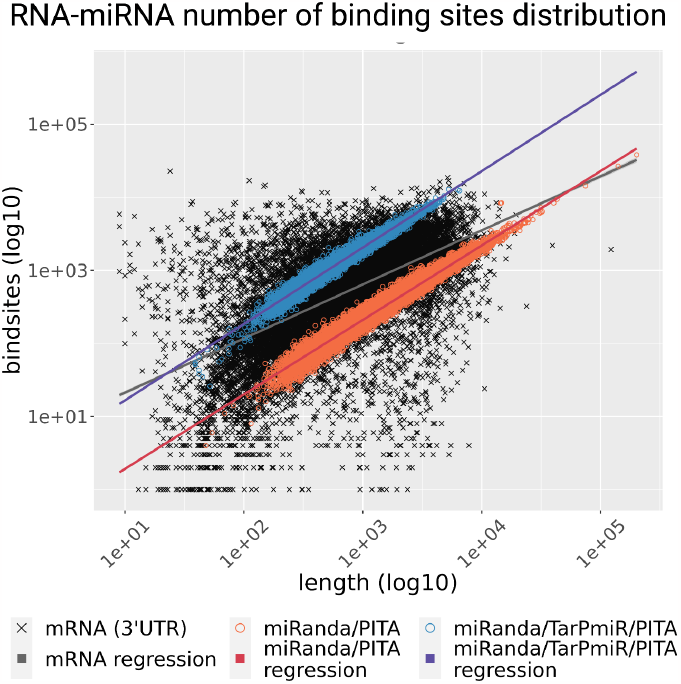
Number of miRNA binding sites versus transcript length for linear and circular RNA. For the 3’UTRs of mRNAs, the number of binding sites was inferred from miRWalk 3.0. circRNA-miRNA binding sites were counted if they were reported by two of the three prediction methods employed, i.e., miRanda, TarPmiR, and PITA. circRNAs form two clusters that can be explained by the different target site prediction methods used. Linear regression models were fit to each of the groups to show the trend of the association.

Compared to the prediction in 3’ UTRs (also based on TarPmiR), circRNAs show a comparably high ratio between the number of binding sites and the length. We observed two distinct circRNA clusters despite employing the same three miRNA binding site prediction methods (miRanda, TarPmiR, PITA) for all of them. We only accept a miRNA binding site for a circRNA if it was predicted by at least two prediction methods. It appears that all three miRNA binding site prediction tools were able to identify miRNA binding sites in the circRNAs of the blue cluster, while only miRanda and PITA, but not TarPmiR, could predict miRNA binding sites of the circRNAs in the red cluster. This observation was not expected as TarPmiR, in general, predicts overall more binding sites than miRanda or PITA. An interesting question is hence if TarPmiR is able to differentiate between circRNAs that are active miRNA sponges and those that have other functions. Previous research has defined three types of circular RNAs based on structural features - exonic circular (ecirc) RNAs, circular intronic RNA (ciRNA), and exon-intron circRNA (EIciRNA) [53–55]. It has been suggested that ecircRNAs function predominantly through a miRNA sponging effect in the cytoplasm, whereas other circular RNA forms (e.g., ciRNA and ElciRNA) function in the nucleus to regulate gene transcription [56–58]. Hence, circRNAs that are functional in the nucleus could have fewer miRNA binding sites. To test alternative explanations, we checked if clusters differed by (1) biotype of the circRNA host gene (i.e., coding or non-coding gene, Suppl. Fig. 12a), (2) genesis, i.e., the splicing method of the circRNA (ElciRNA, ciRNA, ecircRNA, Suppl. Fig. 12b [35]), or (3) circRNA expression level (Suppl. Fig. 12c). The observed clusters did not differ in any of these categories, and further work is needed to elucidate if these results are related to other structural features. It should also be noted that TarPmiR was not trained specifically on circRNAs and that a prediction method tailored towards circRNAs should be developed when suitable experimental data become available. Additionally, we investigated the relationship between the number of miRNA binding sites and the SPONGE [12] correlation scores associated with each circRNA and found that these scores seem to have no apparent correlation to the number of miRNA binding sites, although a very large number of binding sites seems to be advantageous for generating more elevated scores (i.e., over 0.5) in comparison to circRNAs with lower miRNA binding potential, that are only rarely able to reach comparably high correlation values (Suppl. Fig. 13a-b).

## CONCLUSION AND OUTLOOK

We developed a new circRNA processing and analysis pipeline consisting of three modules harboring multiple current state-of-the-art methods: 1) circRNA detection, 2) miRNA detection, and 3) detection of sponging events between circRNAs, ceRNAs, and miRNAs. To the best of our knowledge, it is the first comprehensive circRNA pipeline to detect, quantify and annotate circRNAs as well as to determine their sponging activity. The latter allows users to bring circRNAs into a functional context with other RNAs, such as mRNAs and lncRNAs, through a joint ceRNA network which is mediated by miRNA sponging. Wen et al. recently highlighted the need for a further extensive investigation into circRNAs due to their enormous potential to explain human diseases like cardiovascular, autoimmune, and cancers [59]. circRNAs are also known to be involved in brain development, brain cell differentiation, and neuronal signaling [29,49]. To demonstrate the capabilities of the circRNA-sponging pipeline, we hence re-analyzed a public data set where we investigated circRNAs across different mouse brain regions. Using our pipeline, we could offer novel insights into circRNA biology across tissues of the brain. We showed that differences in circRNA splicing could be revealed when considering the expression of circRNAs relative to the expression of a host gene, similar to how alternative splicing events are detected by considering exon or intron inclusion. Our pipeline is the first to routinely incorporate differential splicing analysis between linear and circular transcripts of the same genes, allowing to better differentiate between changes in expression and changes in splicing. We further placed our findings into the context of miRNA sponging, demonstrating that circRNA exerts regulatory control over a vast number of transcripts. Finally, we showed that the number of binding sites in circRNAs correlated well with their length and observed that TarPmiR’s machine-learning strategy identifies a subset of circRNAs that could indicate promising candidates for miRNA sponging. Further work is needed to investigate if these two classes represent structurally different circRNAs, such as ecircRNAs, ciRNAs, or ElciRNAs, or if this observation can be explained by differences in the miRNA prediction methods with no biological implication at all.

In the future, we plan to extend the circRNA-sponging pipeline with additional features. As various functions other than miRNA sponging have been attributed to circRNAs [60], we see room for expanding the features towards, e.g., investigating the protein-coding potential of circRNAs [1]. We further seek to integrate circRNA-sponging into ongoing community efforts such as nf-core [30] to build up long-term support for maintaining and expanding this pipeline. In summary, the circRNA-sponging pipeline is a powerful tool to detect, investigate, and analyze circRNAs and their sponging effects and thus, it helps researchers consider circRNAs as a routine aspect in RNA-seq and miRNA-seq data analysis.

## Supporting information

Supplementary Data is available at biorxiv online.

## AVAILABILITY

Data are available in GEO under GSE100265 (rRNA-depleted RNA-seq) and GSE93129 (miRNA).

circRNA-sponging is available under the GPL v.3.0 license at: https://github.com/biomedbigdata/circRNA-sponging Dockerhub image is available under:

https://hub.docker.com/repository/docker/leonschwartzdocker/circrnapipeline/general The results presented in this manuscript are available as RData objects at: https://doi.org/10.6084/m9.figshare.21864948

## ACKNOWLEDGMENTS

We thank Bahar Tercan, Mauro A. A. Castro, A. Gordon Robertson, Katherine A. Hoadley, and Victoria K. Cortessis for the fruitful discussions. We thank Christina Trummer for the logo design. Figures were created with https://www.biorender.com. Parts of the images were designed using resources from Flaticon.com.

## REFERENCES

1. Miao Q, Ni B, Tang J. Coding potential of circRNAs: new discoveries and challenges. PeerJ. 2021;9: e10718.

2. Yu C-Y, Kuo H-C. The emerging roles and functions of circular RNAs and their generation. J Biomed Sci. 2019;26: 29.

3. Jeck WR, Sharpless NE. Detecting and characterizing circular RNAs. Nat Biotechnol. 2014;32: 453–461.

4. Lasda E, Parker R. Circular RNAs: diversity of form and function. RNA. 2014;20: 1829–1842.

5. Suzuki H, Zuo Y, Wang J, Zhang MQ, Malhotra A, Mayeda A. Characterization of RNase R-digested cellular RNA source that consists of lariat and circular RNAs from pre-mRNA splicing. Nucleic Acids Res. 2006;34: e63.

6. Mester-Tonczar J, Hašimbegović E, Spannbauer A, Traxler D, Kastner N, Zlabinger K, et al. Circular RNAs in Cardiac Regeneration: Cardiac Cell Proliferation, Differentiation, Survival, and Reprogramming. Front Physiol. 2020;11: 580465.

7. Szabo L, Salzman J. Detecting circular RNAs: bioinformatic and experimental challenges. Nat Rev Genet. 2016;17: 679–692.

8. Jeck WR, Sorrentino JA, Wang K, Slevin MK, Burd CE, Liu J, et al. Circular RNAs are abundant, conserved, and associated with ALU repeats. RNA. 2013;19: 141–157.

9. Zhang Z, Yang T, Xiao J. Circular RNAs: Promising Biomarkers for Human Diseases. EBioMedicine. 2018;34: 267–274.

10. Kristensen LS, Hansen TB, Venø MT, Kjems J. Circular RNAs in cancer: opportunities and challenges in the field. Oncogene. 2018;37: 555–565.

11. Qu S, Yang X, Li X, Wang J, Gao Y, Shang R, et al. Circular RNA: A new star of noncoding RNAs. Cancer Lett. 2015;365: 141–148.

12. List M, Dehghani Amirabad A, Kostka D, Schulz MH. Large-scale inference of competing endogenous RNA networks with sparse partial correlation. Bioinformatics. 2019;35: i596–i604.

13. Hoffmann M, Pachl E, Hartung M, Stiegler V. SPONGEdb: a pan-cancer resource for competing endogenous RNA interactions. Narodonaselenie. 2021. Available: https://academic.oup.com/narcancer/article-abstract/3/1/zcaa042/6066568

14. Kartha RV, Subramanian S. Competing endogenous RNAs (ceRNAs): new entrants to the intricacies of gene regulation. Front Genet. 2014;5: 8.

15. Weinstein JN, Collisson EA, Mills GB, Shaw KRM, Ozenberger BA, Ellrott K, et al. The Cancer Genome Atlas Pan-Cancer analysis project. Nat Genet. 2013;45: 1113–1120.

16. Salmena L, Poliseno L, Tay Y, Kats L, Pandolfi PP. A ceRNA hypothesis: the Rosetta Stone of a hidden RNA language? Cell. 2011;146: 353–358.

17. Hansen TB, Jensen TI, Clausen BH, Bramsen JB, Finsen B, Damgaard CK, et al. Natural RNA circles function as efficient microRNA sponges. Nature. 2013;495: 384–388.

18. Li M, Ding W, Tariq MA, Chang W, Zhang X, Xu W, et al. A circular transcript of ncx1 gene mediates ischemic myocardial injury by targeting miR-133a-3p. Theranostics. 2018;8: 5855–5869.

19. Yu KH-O, Shi CH, Wang B, Chow SH-C, Chung GT-Y, Lung RW-M, et al. Quantifying full-length circular RNAs in cancer. Genome Res. 2021. doi:10.1101/gr.275348.121

20. Chen L, Wang C, Sun H, Wang J, Liang Y, Wang Y, et al. The bioinformatics toolbox for circRNA discovery and analysis. Brief Bioinform. 2021;22: 1706–1728.

21. Zhang X-O, Dong R, Zhang Y, Zhang J-L, Luo Z, Zhang J, et al. Diverse alternative back-splicing and alternative splicing landscape of circular RNAs. Genome Res. 2016;26: 1277–1287.

22. Zhang X-O, Wang H-B, Zhang Y, Lu X, Chen L-L, Yang L. Complementary sequence-mediated exon circularization. Cell. 2014;159: 134–147.

23. Memczak S, Jens M, Elefsinioti A, Torti F, Krueger J, Rybak A, et al. Circular RNAs are a large class of animal RNAs with regulatory potency. Nature. 2013;495: 333–338.

24. Meng X, Chen Q, Zhang P, Chen M. CircPro: an integrated tool for the identification of circRNAs with protein-coding potential. Bioinformatics. 2017;33: 3314–3316.

25. Szabo L, Morey R, Palpant NJ, Wang PL, Afari N, Jiang C, et al. Statistically based splicing detection reveals neural enrichment and tissue-specific induction of circular RNA during human fetal development. Genome Biol. 2015;16: 126.

26. Westholm JO, Miura P, Olson S, Shenker S, Joseph B, Sanfilippo P, et al. Genome-wide analysis of drosophila circular RNAs reveals their structural and sequence properties and age-dependent neural accumulation. Cell Rep. 2014;9: 1966–1980.

27. Glaźar P, Papavasileiou P, Rajewsky N. circBase: a database for circular RNAs. RNA. 2014;20: 1666–1670.

28. Dong R, Ma X-K, Li G-W, Yang L. CIRCpedia v2: An Updated Database for Comprehensive Circular RNA Annotation and Expression Comparison. Genomics Proteomics Bioinformatics. 2018;16: 226–233.

29. Piwecka M, Glažar P, Hernandez-Miranda LR, Memczak S, Wolf SA, Rybak-Wolf A, et al. Loss of a mammalian circular RNA locus causes miRNA deregulation and affects brain function. Science. 2017;357. doi:10.1126/science.aam8526

30. Ewels PA, Peltzer A, Fillinger S, Patel H, Alneberg J, Wilm A, et al. The nf-core framework for community-curated bioinformatics pipelines. Nat Biotechnol. 2020;38: 276–278.

31. Dobin CA, Davis FS. J. Drenkow, C. Zaleski, S. Jha, P. Batut, M. Chaisson. and TR Gingeras. 2013. STAR: ultrafast universal RNA-seq aligner 舰; 2013.

32. Frazee AC, Jaffe AE, Langmead B, Leek JT. Polyester: simulating RNA-seq datasets with differential transcript expression. Bioinformatics. 2015;31: 2778–2784.

33. Love MI, Huber W, Anders S. Moderated estimation of fold change and dispersion for RNA-seq data with DESeq2. Genome Biol. 2014;15: 550.

34. Bray NL, Pimentel H, Melsted P, Pachter L. Erratum: Near-optimal probabilistic RNA-seq quantification. Nat Biotechnol. 2016;34: 888.

35. Trincado JL, Entizne JC, Hysenaj G, Singh B, Skalic M, Elliott DJ, et al. SUPPA2: fast, accurate, and uncertainty-aware differential splicing analysis across multiple conditions. Genome Biol. 2018;19: 40.

36. Enright AJ, John B, Gaul U, Tuschl T, Sander C, Marks DS. MicroRNA targets in Drosophila. Genome Biol. 2003;5: R1.

37. Kertesz M, Iovino N, Unnerstall U, Gaul U, Segal E. The role of site accessibility in microRNA target recognition. Nat Genet. 2007;39: 1278–1284.

38. Ding J, Li X, Hu H. TarPmiR: a new approach for microRNA target site prediction. Bioinformatics. 2016;32: 2768–2775.

39. Min H, Yoon S. Got target? Computational methods for microRNA target prediction and their extension. Exp Mol Med. 2010;42: 233–244.

40. Karagkouni D, Paraskevopoulou MD, Tastsoglou S, Skoufos G, Karavangeli A, Pierros V, et al. DIANA-LncBase v3: indexing experimentally supported miRNA targets on non-coding transcripts. Nucleic Acids Res. 2020;48: D101–D110.

41. Hsu S-D, Lin F-M, Wu W-Y, Liang C, Huang W-C, Chan W-L, et al. miRTarBase: a database curates experimentally validated microRNA–target interactions. Nucleic Acids Research. 2011. pp. D163–D169. doi:10.1093/nar/gkq1107

42. Huang H-Y, Lin Y-C-D, Li J, Huang K-Y, Shrestha S, Hong H-C, et al. miRTarBase 2020: updates to the experimentally validated microRNA-target interaction database. Nucleic Acids Res. 2020;48: D148–D154.

43. Sticht C, De La Torre C, Parveen A, Gretz N. miRWalk: An online resource for prediction of microRNA binding sites. PLoS One. 2018;13: e0206239.

44. Friedländer MR, Mackowiak SD, Li N, Chen W, Rajewsky N. miRDeep2 accurately identifies known and hundreds of novel microRNA genes in seven animal clades. Nucleic Acids Res. 2012;40: 37–52.

45. Boniolo F, Hoffmann M, Roggendorf N, Tercan B, Baumbach J, Castro MAA, et al. spongEffects: ceRNA modules offer patient-specific insights into the miRNA regulatory landscape. Bioinformatics. 2023. doi:10.1093/bioinformatics/btad276

46. Hänzelmann S, Castelo R, Guinney J. GSVA: gene set variation analysis for microarray and RNA-seq data. BMC Bioinformatics. 2013;14: 7.

47. Jerby-Arnon L, Shah P, Cuoco MS, Rodman C, Su M-J, Melms JC, et al. A Cancer Cell Program Promotes T Cell Exclusion and Resistance to Checkpoint Blockade. Cell. 2018;175: 984–997.e24.

48. Kuhn M. Building Predictive Models in R Using the caret Package. J Stat Softw. 2008;28: 1–26.

49. Hanan M, Soreq H, Kadener S. CircRNAs in the brain. RNA Biol. 2017;14: 1028–1034.

50. Rybak-Wolf A, Stottmeister C, Glažar P, Jens M, Pino N, Giusti S, et al. Circular RNAs in the Mammalian Brain Are Highly Abundant, Conserved, and Dynamically Expressed. Mol Cell. 2015;58: 870–885.

51. Galea JM, Vazquez A, Pasricha N, de Xivry J-JO, Celnik P. Dissociating the roles of the cerebellum and motor cortex during adaptive learning: the motor cortex retains what the cerebellum learns. Cereb Cortex. 2011;21: 1761–1770.

52. Jiang C, Zeng X, Shan R, Wen W, Li J, Tan J, et al. The Emerging Picture of the Roles of CircRNA-CDR1as in Cancer. Front Cell Dev Biol. 2020;8: 590478.

53. Yang L, Fu J, Zhou Y. Circular RNAs and Their Emerging Roles in Immune Regulation. Front Immunol. 2018;9: 2977.

54. Xiao J, Joseph S, Xia M, Teng F, Chen X, Huang R, et al. Circular RNAs Acting as miRNAs’ Sponges and Their Roles in Stem Cells. J Clin Med Res. 2022;11. doi:10.3390/jcm11102909

55. Zhang Y, Tian Z, Ye H, Sun X, Zhang H, Sun Y, et al. Emerging functions of circular RNA in the regulation of adipocyte metabolism and obesity. Cell Death Discov. 2022;8: 268.

56. Zhang Y, Zhang X-O, Chen T, Xiang J-F, Yin Q-F, Xing Y-H, et al. Circular intronic long noncoding RNAs. Mol Cell. 2013;51: 792–806.

57. Li Z, Huang C, Bao C, Chen L, Lin M, Wang X, et al. Exon-intron circular RNAs regulate transcription in the nucleus. Nat Struct Mol Biol. 2015;22: 256–264.

58. Li Z, Huang C, Bao C, Chen L, Lin M, Wang X, et al. Corrigendum: Exon-intron circular RNAs regulate transcription in the nucleus. Nat Struct Mol Biol. 2017;24: 194.

59. Wen G, Zhou T, Gu W. The potential of using blood circular RNA as liquid biopsy biomarker for human diseases. Protein Cell. 2021;12: 911–946.

60. Nielsen AF, Bindereif A, Bozzoni I, Hanan M, Hansen TB, Irimia M, et al. Best practice standards for circular RNA research. Nat Methods. 2022;19: 1208–1220.

