## Supplementary Data is available at biorxiv online. for "circRNA-sponging: a pipeline for extensive analysis of circRNA expression and their role in miRNA sponging"

#### Supplementary Figure 1: circRNA detection

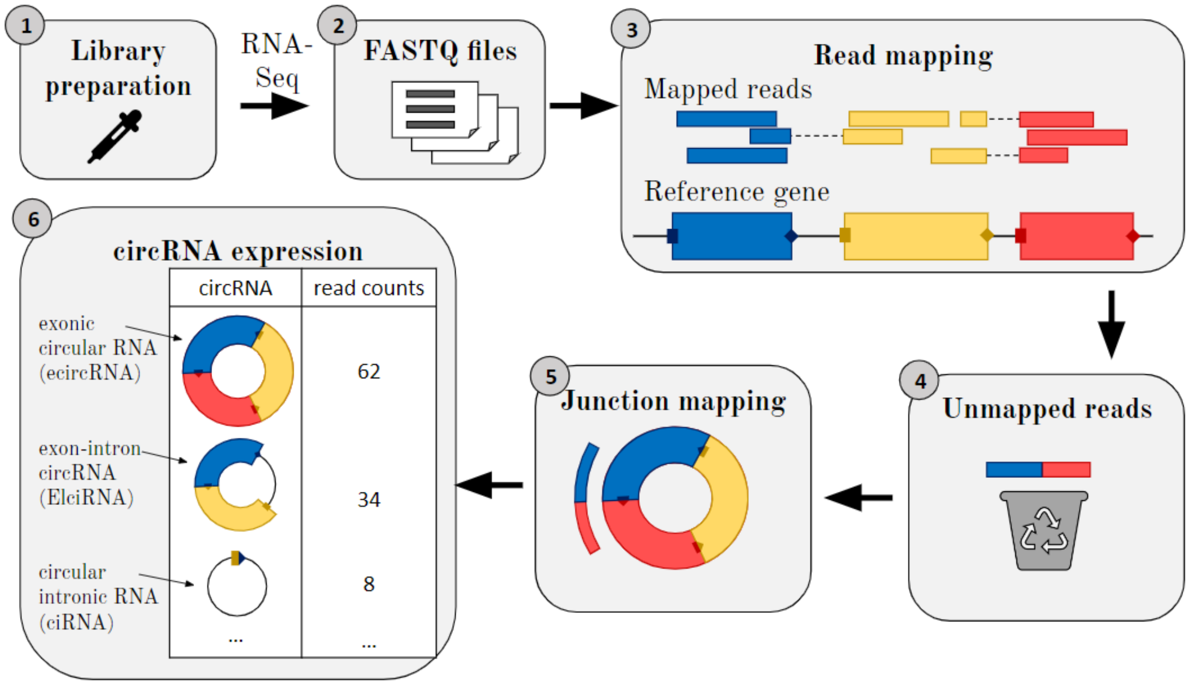


**Supplementary Figure 1:** First, data has to be prepared in the laboratory by library preparation and RNA-sequencing (e.g., totalRNA-seq). Then the next step is to map the RNA sequencing reads to a reference genome or transcriptome. This is typically done using alignment tools such as TopHat, HISAT2, STAR, or Bowtie2. Then backsplice junctions have to be identified: In this step, the mapped reads are analyzed to identify backsplice junctions, which are the junctions formed when the 5' and 3' ends of a linear RNA molecule are covalently linked to form a circular RNA. Several tools, such as CIRCexplorer, find_circ, and circRNA_finder, can be used to identify backsplice junctions. Filtering: In this step, low-quality and false-positive backsplice junctions are filtered out based on several criteria, such as read coverage, number of supporting reads, and mapping quality. Then we obtain the circRNA expression that includes exonic circular (ecirc) RNAs, circular intronic RNA (ciRNA), and exon-intron circRNA (EIciRNA).

### Supplementary Figure 2: evaluation of existing methods to detect circRNAs and miRNA binding sites

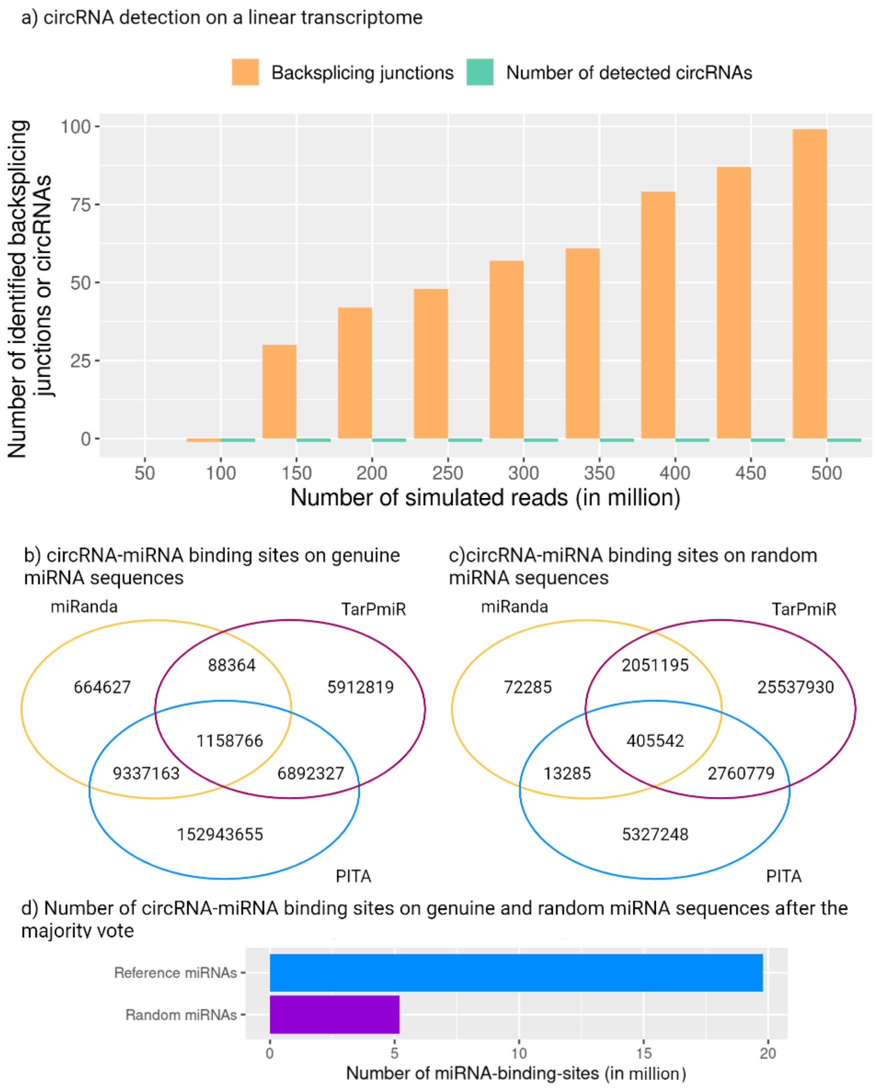


**Supplementary Figure 2**: a) We used polyester [1] to simulate linear reads with sequencing depths between 100 and 500 million reads from a linear mouse transcriptome (GRCm38). We used the circRNA-sponging pipeline to detect circRNAs and detect very few false-positive backsplicing junctions (0-100, slowly increasing with increasing sequencing depth) in the linear transcript, however, we can not detect circRNAs (i.e., a false-discovery rate of 0%). b-d) We randomly sampled 672 miRNAs (i.e., the equal number of miRNAs that were used in the majority vote of the mouse brain dataset). The artificial miRNA sequences were generated from randomly sampled DNA bases with lengths between 19 and 23 nucleotides, as suggested in [2]. We then predicted miRNA binding sites between the correct circRNAs and the randomly sampled miRNAs with miRanda [3], PITA [4], and TarPmiR [5]. We can see that some miRNA binding sites are predicted in (c), but in total, up to 97% fewer miRNA binding sites in some of the methods than in the genuine run (b) of the majority vote. (d) shows that, in total, we find significantly fewer miRNA binding sites with randomly generated miRNAs.

### Supplementary Figure 3: Annotation and differential overall circRNA expression in tissues

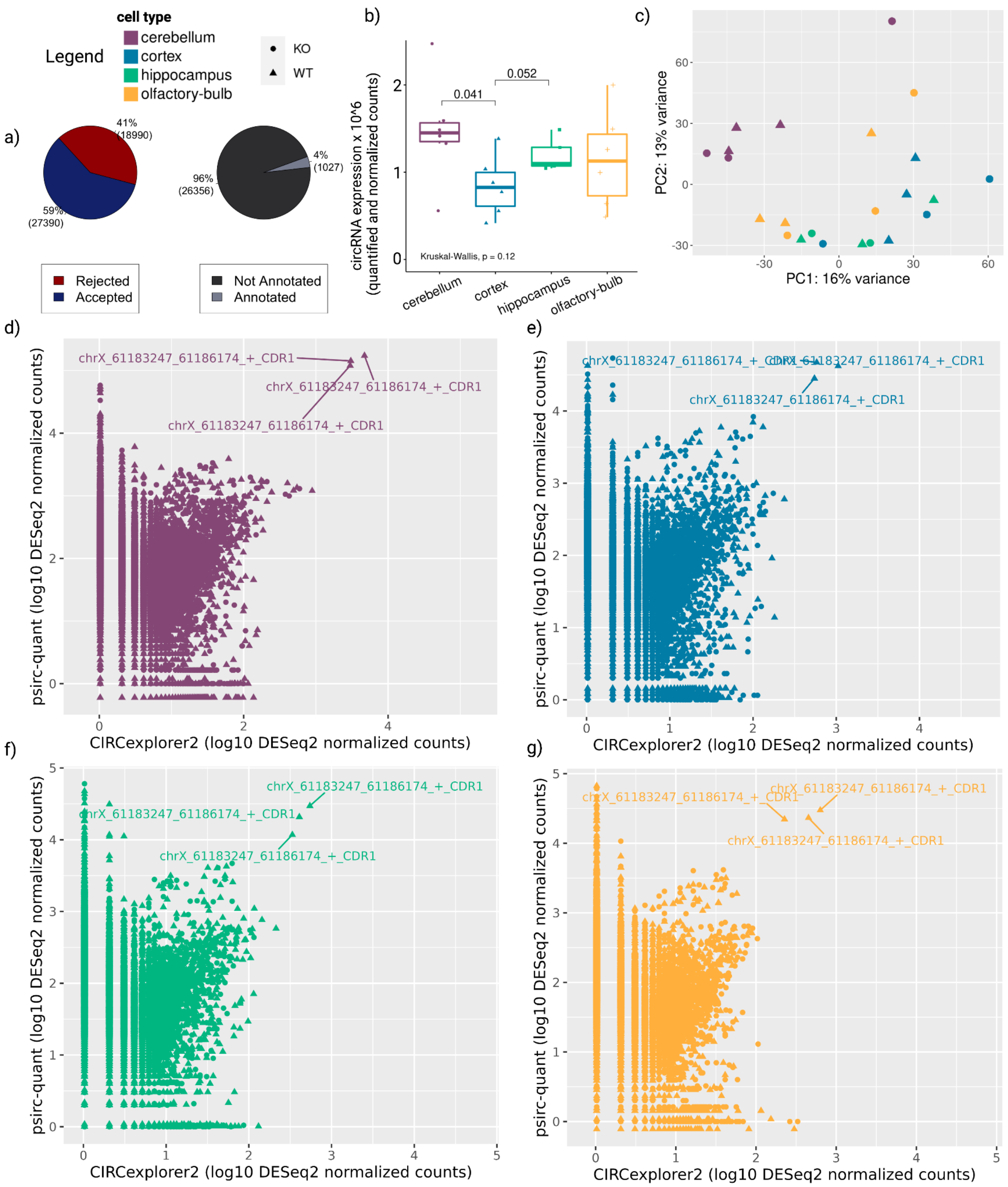


**Supplementary Figure 3:** a) Proportion of circRNA that have passed the filters that were applied by the circRNA sponging pipeline and proportion of circRNAs that we can annotate. b) Expression levels across all samples per tissue and detected circRNAs. c) PCA of all detected circRNAs per tissue. d)-g) differences in circRNA expression between raw counts identified by CIRCexplorer2 and quantified counts by psirc.

#### Supplementary Figure 4: Correlation distribution between circRNA-miRNA pairs that have (a) no common binding sites and (b) that have at least 10 common binding sites

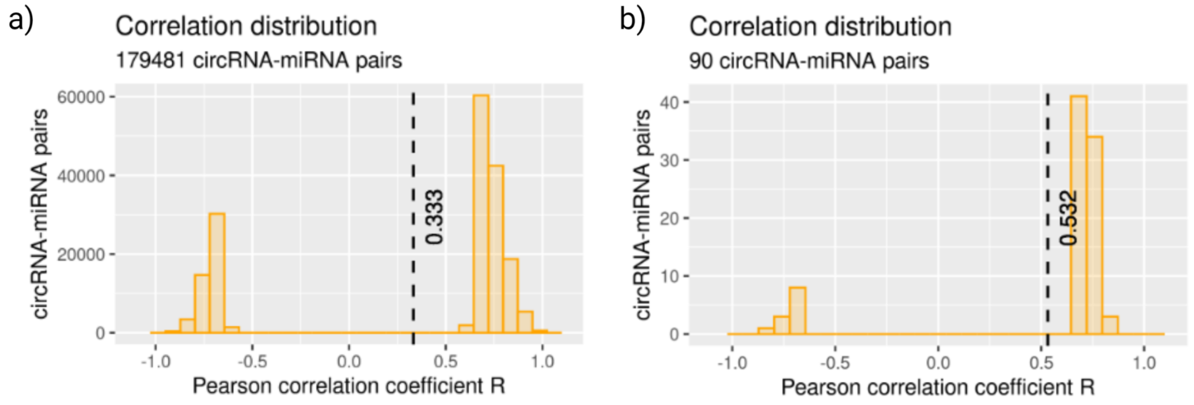


**Supplementary Figure 4:** a) Pearson correlation coefficient distribution with no common binding sites between circRNA and miRNAs. b) Pearson correlation coefficient distribution with at least 10 common binding sites.

#### Supplementary Figure 5: PCAs and Heatmaps of known circRNAs

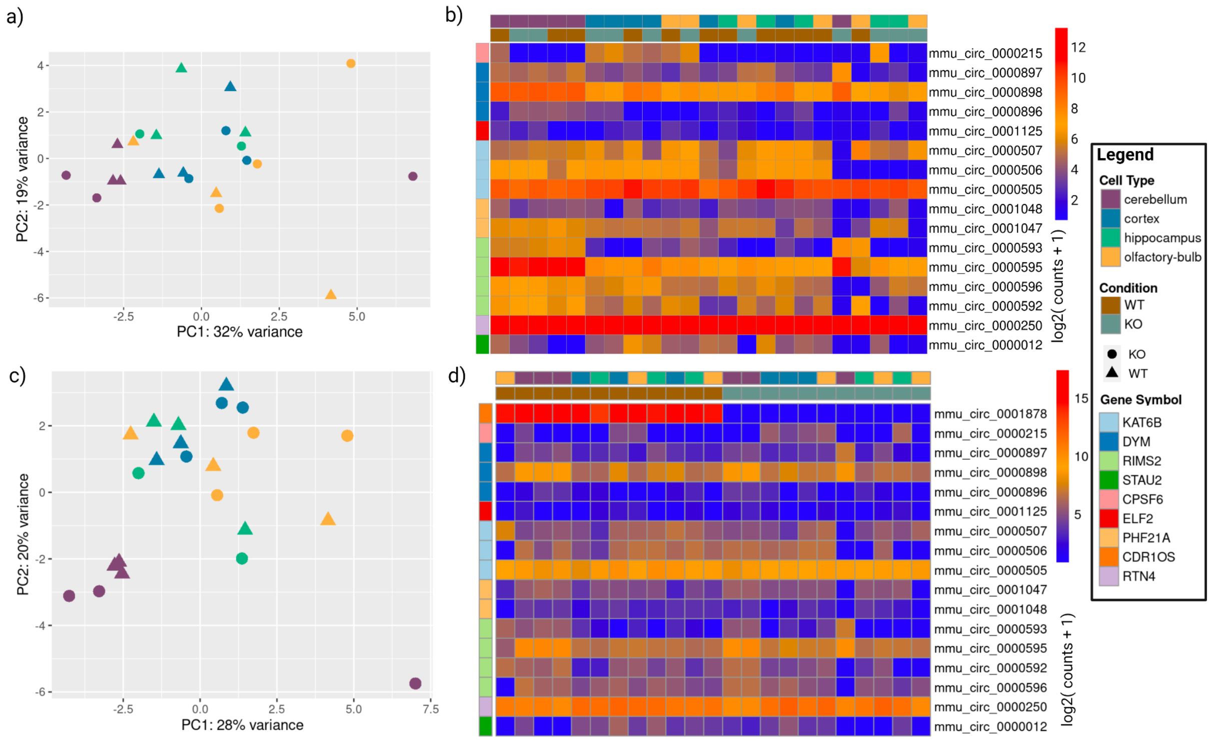


**Supplementary Figure 5:** a) PCA of the circRNAs found by Rybak-Wolf et al. but with CDR1as excluded due to successful targeting in the knockout colored by brain region and symbol by condition. b) Heatmap of the circRNAs found by Rybak-Wolf et al. but with CDR1as excluded due to successful targeting in the knockout colored by brain region and symbol by condition. c) PCA of the circRNAs found by Rybak-Wolf et al. colored by brain region and symbol by condition. d) Heatmap of the circRNAs found by Rybak-Wolf et al. colored by brain region and symbol by condition [6].

### Supplementary Figure 6: circRNA versus host gene expression of known circRNAs

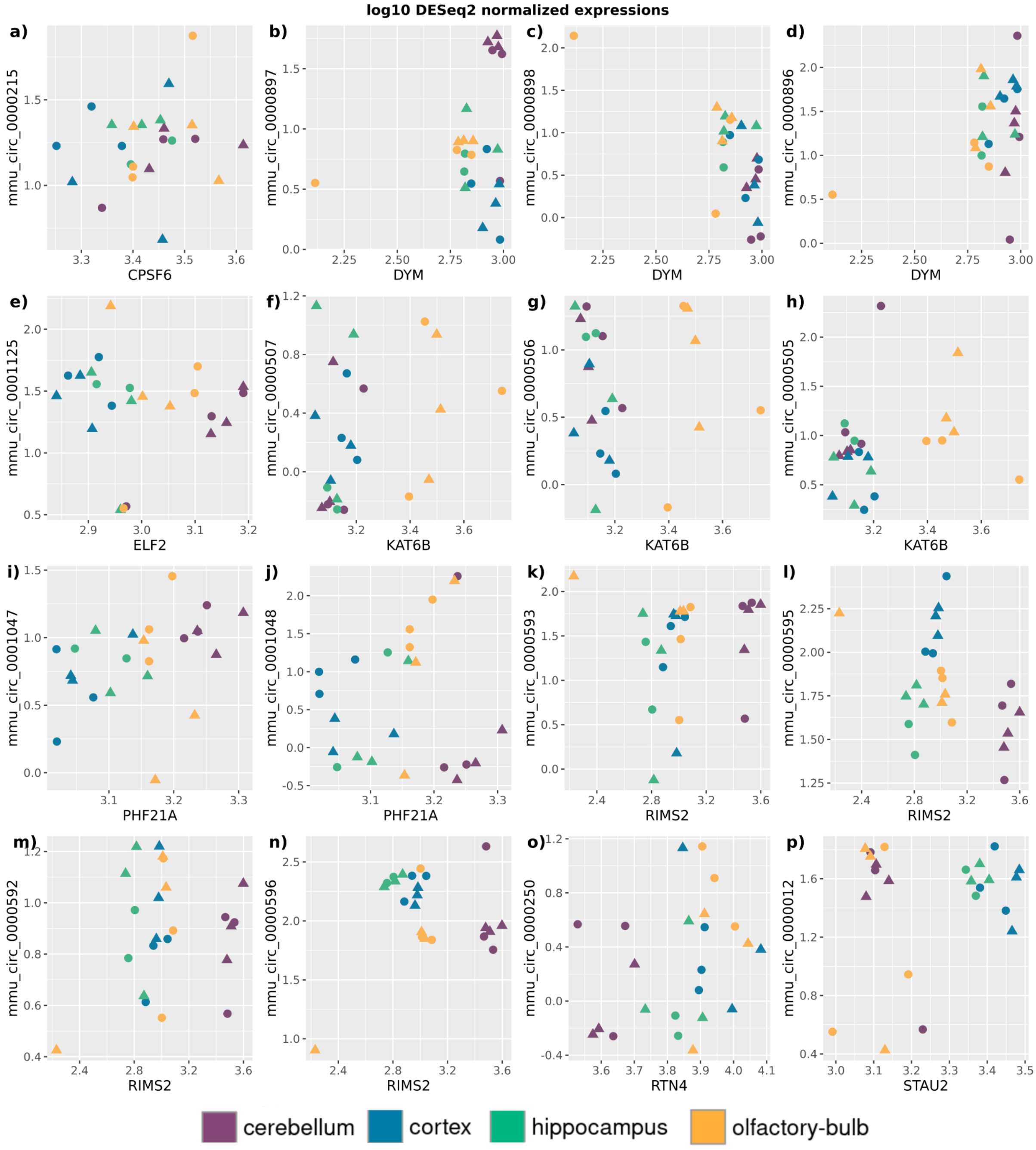


**Supplementary Figure 6:** log10 DESeq2 normalized expression between circRNA and their host gene for circRNAs found by Rybak-Wolf et al. [6].

### Supplementary Figure 7: Linear and circular transcript splicing per host gene

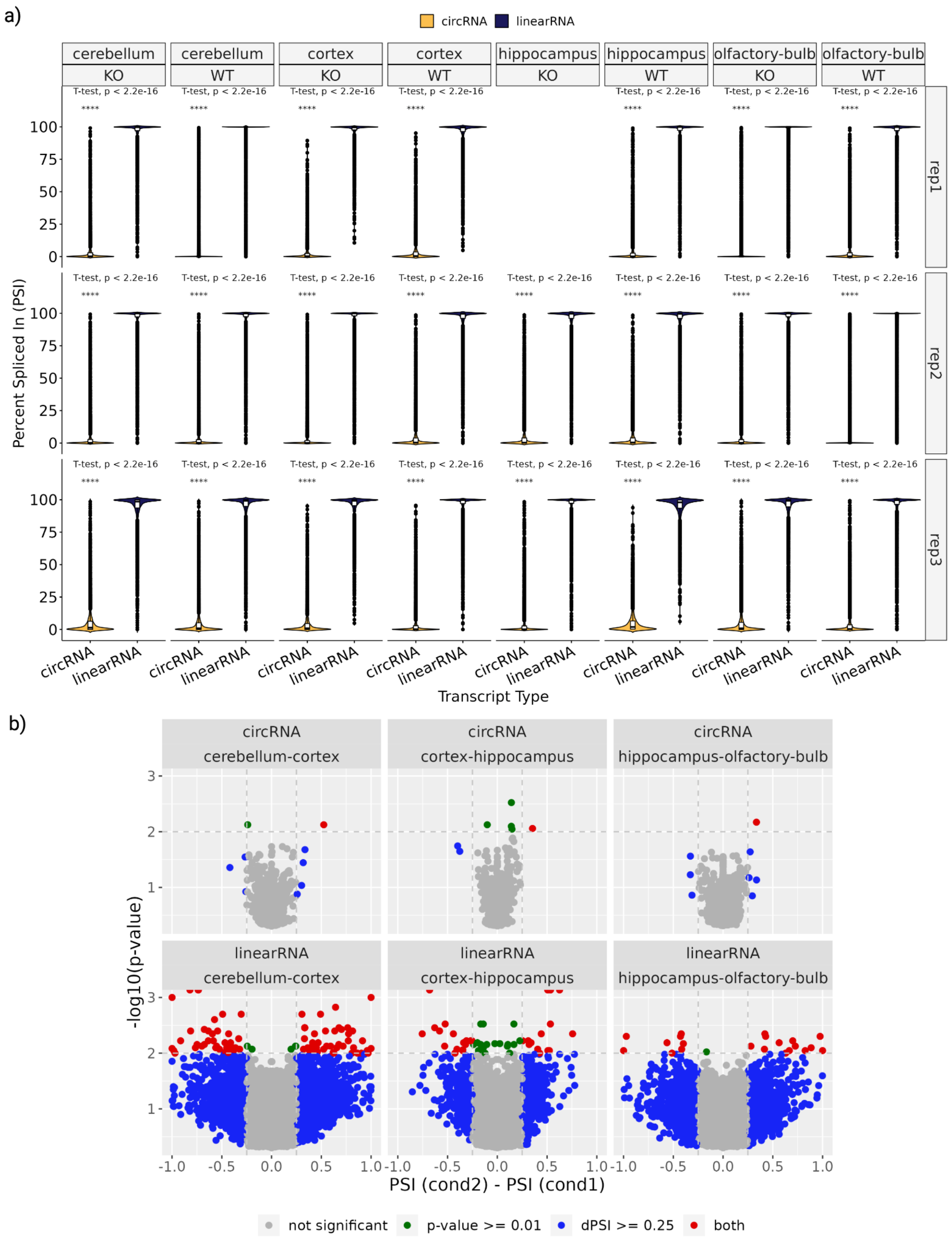


**Supplementary Figure 7:** a) Comparison of percent spliced-in (PSI) values per cell type for linear and circular transcripts within a gene. b) Volcano plots of SUPPA2 differential splicing PSI values between conditions over the negative logarithm of the according p-value for both linear and circular isoforms. Grey points represent non-significant splicing events, green points correspond to significant p-values below 0.01, blue points show a difference in percent spliced-in (PSI) values greater than 25% between conditions, and isoforms marked in red satisfy both filtering criteria. Concerning circular isoforms, the significant circRNAs (shown in red) are *chr7:114319452-114321251_-*, *chr15:34600014-34625031_-,* and *chr8:110298074-110334816_+* for *cerebellum-cortex, cortex-hippocampus,* and *hippocampus-olfactory-bulb,* respectively, which are associated to the host genes of 4933406I18Rik, Hydin, and Nipal2.

### Supplementary Figure 8: Normalized linear and circular transcript splicing per host gene

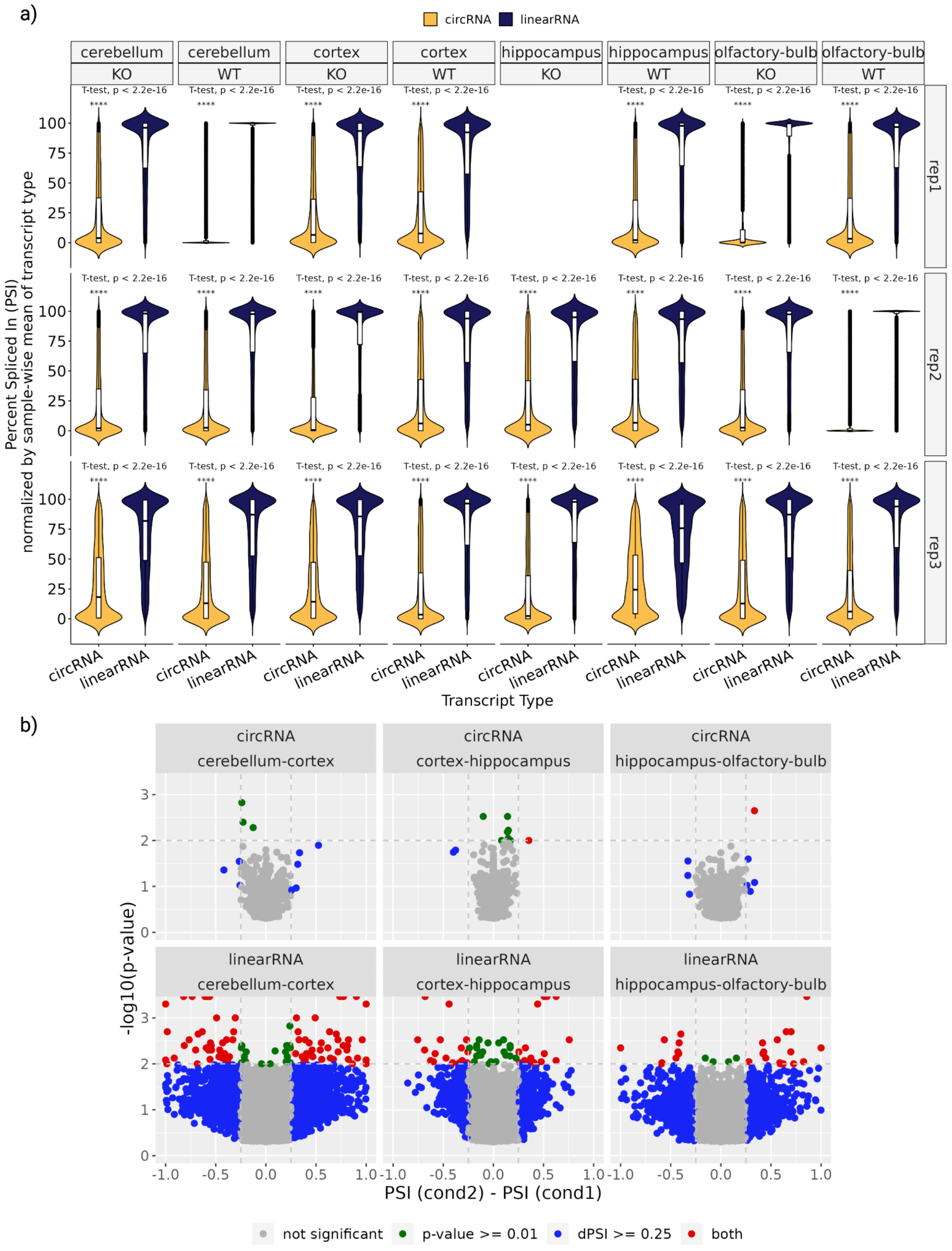


**Supplementary Figure 8:** a) Comparison of percent spliced-in (PSI) values per cell type for linear and circular transcripts within a gene, normalized by the mean PSI value for each transcript type. b) Volcano plots of SUPPA2 differential splicing PSI values between conditions over the negative logarithm of the according p-value for both linear and circular isoforms. The y-axis depicts the p-values in a negative logarithm, and the x-axis shows the change between the PSI values of the conditions. Grey points represent non-significant splicing events, green points correspond to significant p-values below 0.01, blue points show a difference in percent spliced-in (PSI) values greater than 25% between conditions, and isoforms marked in red satisfy both filtering criteria. Concerning circular isoforms, the significant circRNAs (shown in red) are *chr15:34600014-34625031_-* and *chr8:110298074-110334816_+* for *cortex-hippocampus* and *hippocampus-olfactory-bulb,* respectively, which are associated to the host genes of HYDIN and NIPAL2.

### Supplementary Figure 9: spongEffect model performance

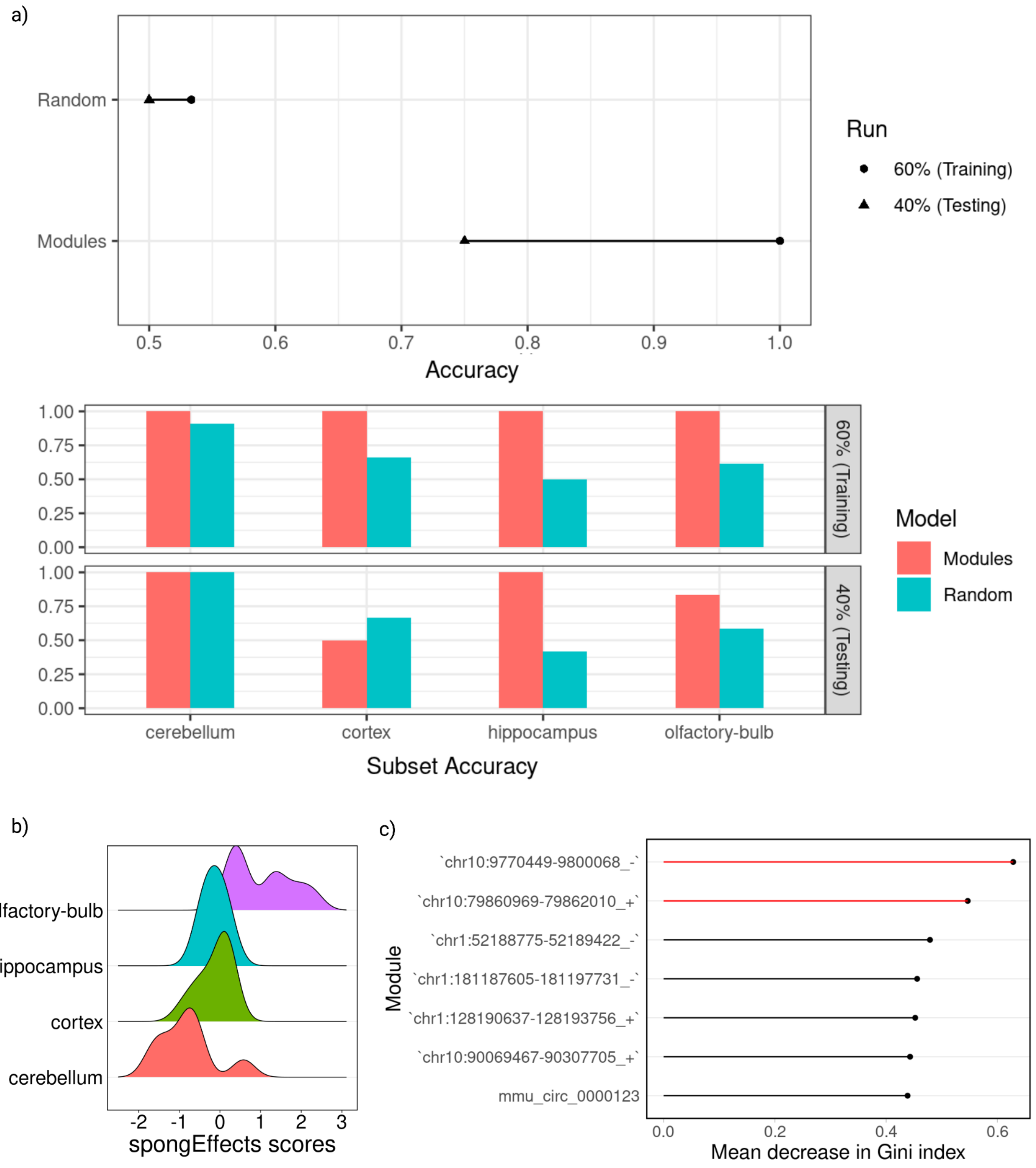


**Supplementary Figure 9:** Evaluation of the spongEffects model performance on the example data. a) Overall prediction model accuracy between randomly generated modules and modules identified by spongEffects and accuracy per brain region between randomly generated modules and modules identified by spongEffects. b) Distribution of the spongEffects enrichment scores per brain region. c) Most predictive modules that were identified by spongEffects were ranked by the decreasing Gini index.

#### Supplementary Figure 10: module expression for central circRNA players

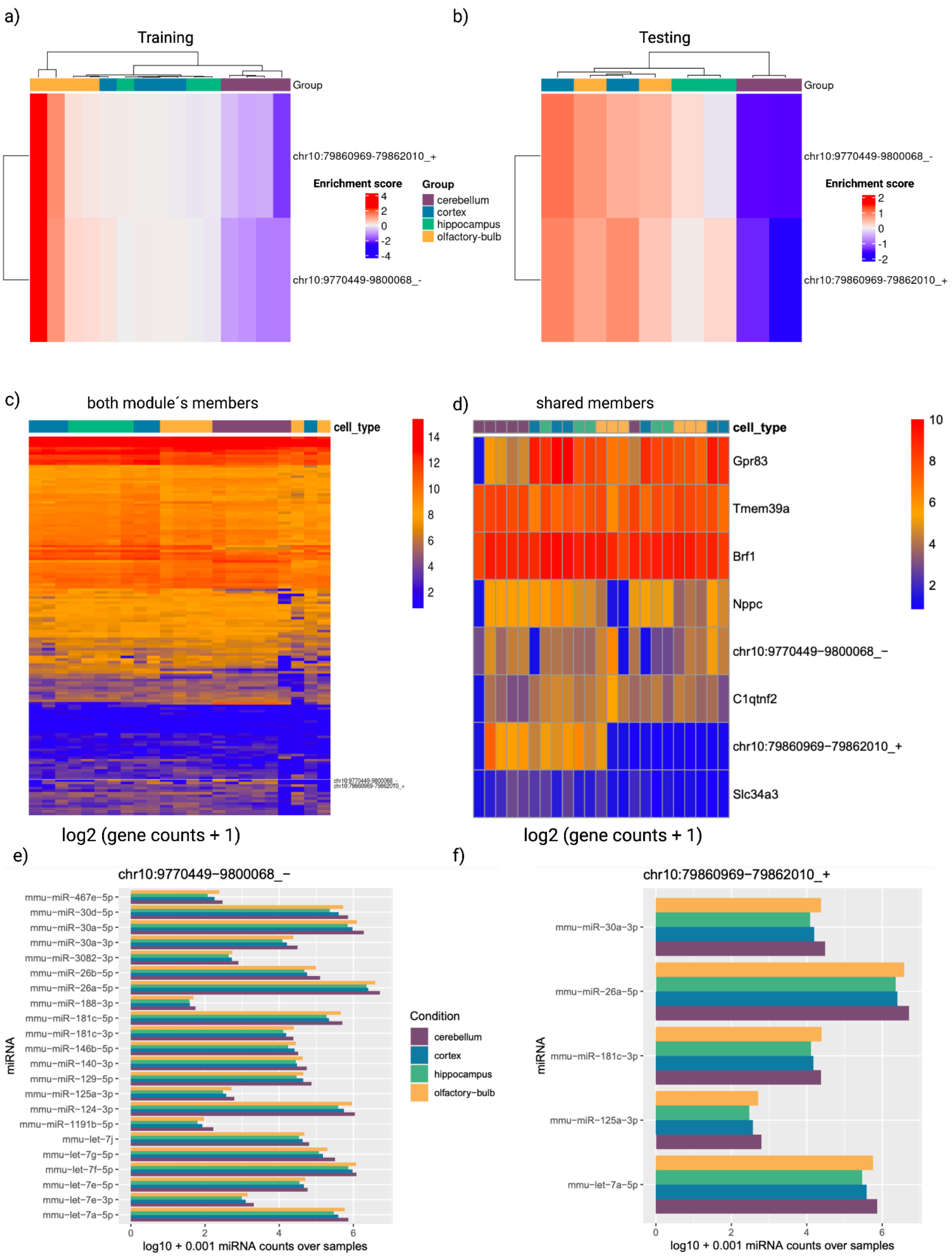


**Supplementary Figure 10:** a)+b) Expression of module members that were associated with circRNAs chr10:9770449-9800068_- and chr10:79860969-79862010_+ in training and test data. c) Heatmap of expression of combined modules members of circRNAs chr10:9770449-9800068_- and chr10:79860969-79862010_+. d) Heatmap of shared modules member between circRNAs chr10:9770449-9800068_- and chr10:79860969-79862010_+. e) Key miRNA players in the ceRNA subnetwork of chr10:9770449-9800068_-. f) Key miRNA players in the ceRNA subnetwork of chr10:79860969-79862010_+.

#### Supplementary Figure 11: module expression for central circRNA players

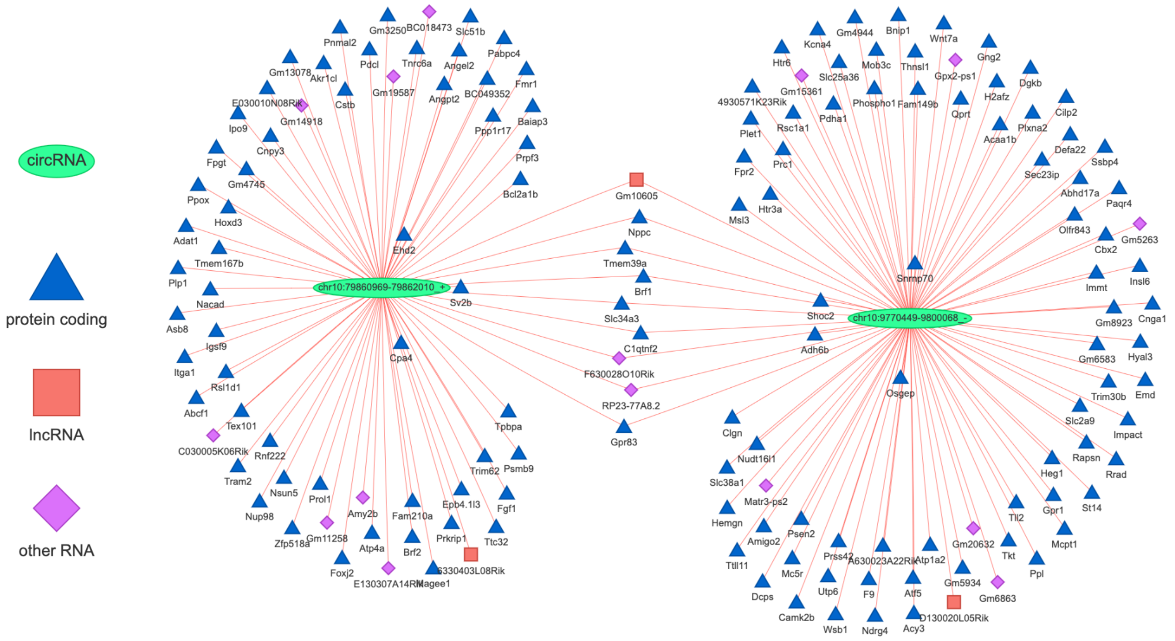


**Supplementary Figure 11:** Visualization of ceRNA subnetworks of chr10:9770449-9800068_- and chr10:79860969-79862010_+ and their overlap.

### Supplementary Figure 12: miRNA binding sites comparison between circRNAs and linear transcripts

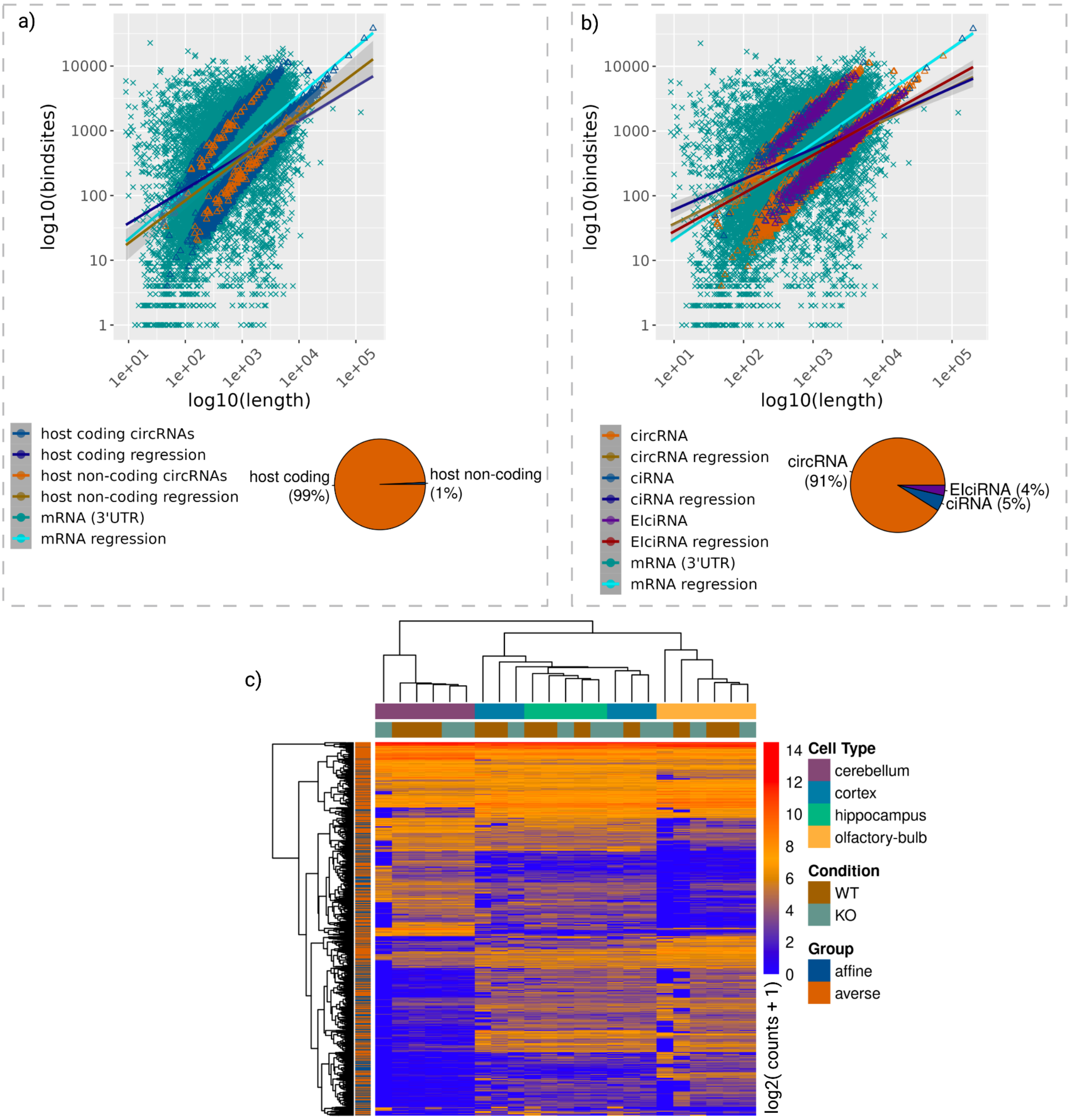


**Supplementary Figure 12:** a+b) Attempt to explain the two circRNA subgroups identified by the number of miRNA binding sites versus transcript length for linear and circular RNA by investigating the biotype of the circRNA host gene (i.e., coding or non-coding gene and exonic, intronic, or exonic-intronic). Next to the legend, this Figure additionally supplies information about the distribution of each circRNA subtype. circRNA types in b) were derived from a combination of CIRCexplorer2 annotate outputs and retained intron splice events generated by SUPPA2 [7]. c) Attempt to explain the two circRNA subgroups identified by the number of miRNA binding sites versus transcript length for linear and circular RNA by investigating the circRNA expression level.

#### Supplementary Figure 13: SPONGE correlation score distributions in relation to the number of miRNA binding sites

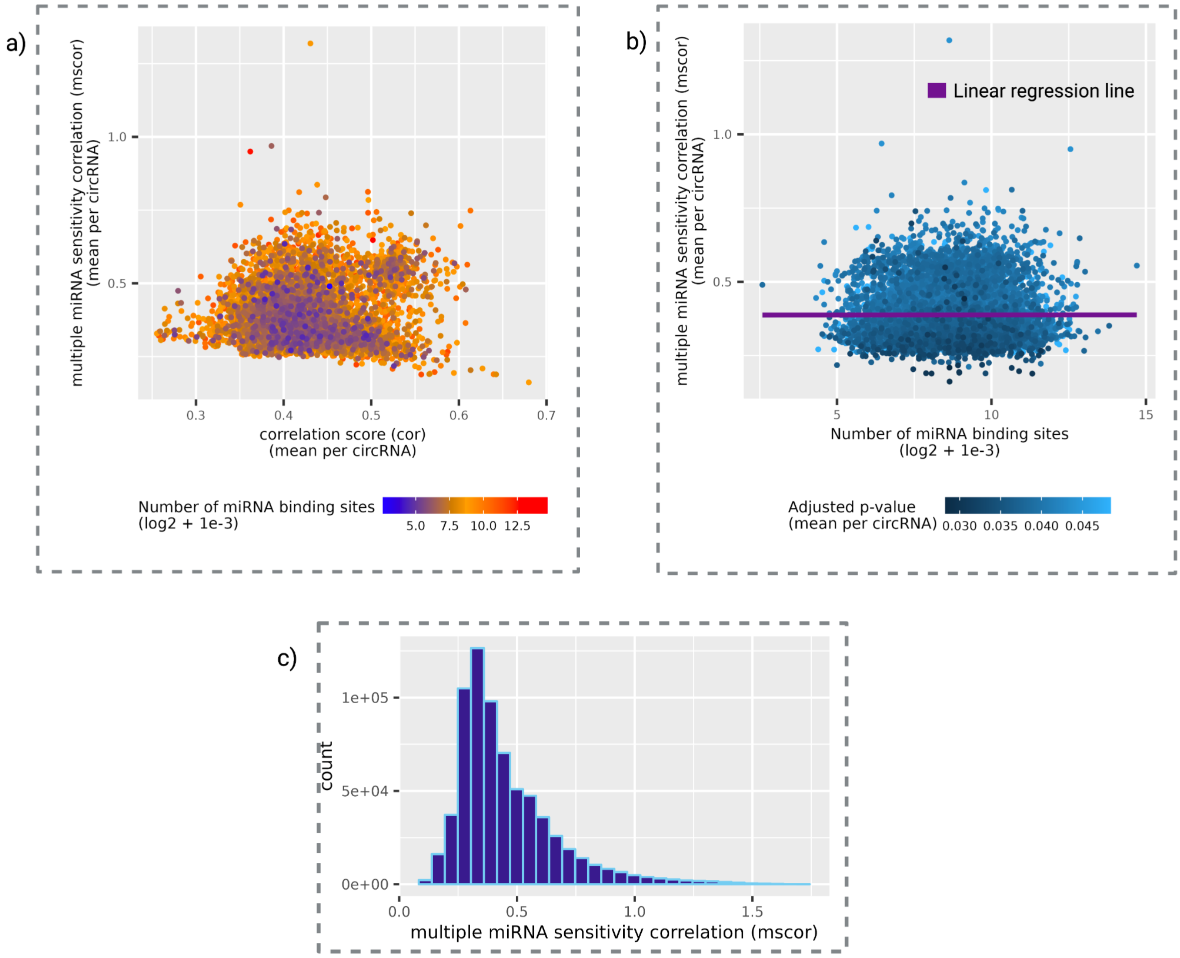


**Supplementary Figure 13:** a) Relation of mean mscor and cor SPONGE scores per circRNA. The color of points represents the number of miRNA binding sites according to circRNA. b) Correlation between mscor and the number of associated miRNA binding sites per circRNA. The violet line shows the linear regression between the two variables. The color of the points shows the mean adjusted p-value per circRNA. c) Distribution of mscor values over the circRNAs.

#### Supplementary Table 1: Number of samples per brain tissue

| variant | cerebellum | | cortex | | hippocampus | | olfactory bulb | | Sum |
|---|---|---|---|---|---|---|---|---|---|
| | WT | KO | WT | KO | WT | KO | WT | KO | |
| sample number | 3 | 3 | 3 | 3 | 3 | 2 | 3 | 3 | 23 |

#### Supplementary Table 3: circRNAs detected for host genes of an experimentally conducted study on circRNAs in different tissues of mouse brains [6]

| Host Gene | circRNA circBase-ID |
|---|---|
| RIMS2 | mmu_circ_0000592<br>mmu_circ_0000593<br>mmu_circ_0000595<br>mmu_circ_0000596 |
| ELF2 | mmu_circ_0001125 |
| PHF21A | mmu_circ_0001047<br>mmu_circ_0001048 |
| MYST4 | mmu_circ_0000505<br>mmu_circ_0000506<br>mmu_circ_0000507 |
| CDR1 | mmu_circ_0001878 |
| STAU2 | mmu_circ_0000012 |
| CPSF6 | mmu_circ_0000215 |
| DYM | mmu_circ_0000896<br>mmu_circ_0000897<br>mmu_circ_0000898 |
| RTN4 | mmu_circ_0000250 |

#### Supplementary Table 3: Highest differentially expressed circRNAs across brain tissues

| Genomic loci | Host gene | (log2fc) cerebellum versus | | |
|---|---|---|---|---|
| | | cortex | hippocampus | olfactory bulb |
| chrX:71369798:71394465:+ | Mtmr1 | 8,04 | 5,28 | 5,2 |
| chr11:87190073:87197015:+ | Trim37 | 7,67 | 5,45 | 5,96 |
| chr10:117077511:117099967:- | Frs2 | 7,64 | 7,43 | 5,71 |
| chr19:56928248:56930261:- | Afap1l2 | 7,03 | 5,36 | 7,81 |
| chr5:122473209:122500948:- | Atp2a2 | 6,78 | 5,05 | 10 |
| chr13:43425029:43436142:- | Ranbp9 | 6,47 | 5,2 | 6,61 |
| chr6:128333832:128337785:- | Tulp3 | 6,32 | 5,8 | 6,72 |
| chr16:21500187:21506821:+ | Vps8 | 6,17 | 6,35 | 6,55 |
| chr6:17660827:17662061:+ | Capza2 | 6,09 | 6,22 | 5,89 |
| chr7:27262167:27268919:+ | Numbl | 5,89 | 6,7 | 7,04 |
| chr6:104642896:104776452:+ | Cntn6 | 5,75 | 5,74 | 5,45 |
| chr5:31470357:31472961:+ | Zfp512 | 5,72 | 5,82 | 5,4 |
| chr4:82320473:82498769:- | Nfib | 5,59 | 6,21 | 5,36 |
| chr5:121273606:121281944:+ | Hectd4 | 5,57 | 5,03 | 7,42 |
| chr3:132734717:132737588:+ | Tbck | 5,54 | 5,14 | 5,55 |
| chr7:111554438:111556529:- | Galnt18 | 5,49 | 5,83 | 6,25 |
| chr11:69363619:69364804:- | Chd3 | 5,49 | 5,61 | 7,62 |
| chr1:36207941:36211486:- | Uggt1 | 5,48 | 5,25 | 7,06 |
| chr15:41800470:41826052:+ | Oxr1 | 5,46 | 5,74 | 5,73 |
| chr11:86298279:86308912:- | Med13 | 5,45 | 5,11 | 6,09 |
| chr1:86159994:86173714:+ | Armc9 | 5,14 | 5,17 | 5,57 |
| chr11:57217771:57229025:+ | Gria1 | 5,13 | 6,56 | 9,09 |
| chr1:139085882:139110485:+ | Dennd1b | 5,04 | 5,6 | 7,01 |
| chr1:63543118:63567968:+ | Adam23 | -5,33 | -6,03 | -5,92 |
| chrX:101546958:101576703:+ | Taf1 | -5,57 | -5,17 | -5,35 |
| chrX:152033698:152047968:+ | Smc1a | -6,02 | -6,73 | -6,43 |

| | | | | |
|---|---|---|---|---|
| chr16:56586928:56606193:+ | Abi3bp | -6,18 | -6,68 | -7,76 |
| chr6:115848905:115850839:- | Mbd4 | -6,25 | -7,36 | -6,34 |
| chr10:82732602:82739265:+ | Hcfc2 | -6,26 | -5,85 | -5,82 |
| chr18:3287903:3327591:- | Crem | -7,41 | -5,16 | -7,34 |
| chr15:101008052:101035705:+ | Scn8a | -7,41 | -8,26 | -5,12 |
| chr9:82876725:82887669:- | Phip | -8,13 | -8,89 | -8,5 |
| chr18:63924811:63935948:+ | Wdr7 | -9,5 | -9,11 | -6,78 |

#### References


1. Frazee AC, Jaffe AE, Langmead B, Leek JT. Polyester: simulating RNA-seq datasets with differential transcript expression. Bioinformatics. 2015;31: 2778–2784.
2. RPubs - generate random DNA sequences. [cited 22 Jun 2023]. Available: https://rpubs.com/oaxacamatt/random_dna_seq
3. Enright AJ, John B, Gaul U, Tuschl T, Sander C, Marks DS. MicroRNA targets in Drosophila. Genome Biol. 2003;5: R1.
4. Kertesz M, Iovino N, Unnerstall U, Gaul U, Segal E. The role of site accessibility in microRNA target recognition. Nat Genet. 2007;39: 1278–1284.
5. Ding J, Li X, Hu H. TarPmiR: a new approach for microRNA target site prediction. Bioinformatics. 2016;32: 2768–2775.
6. Rybak-Wolf A, Stottmeister C, Glažar P, Jens M, Pino N, Giusti S, et al. Circular RNAs in the Mammalian Brain Are Highly Abundant, Conserved, and Dynamically Expressed. Mol Cell. 2015;58: 870–885.
7. Trincado JL, Entizne JC, Hysenaj G, Singh B, Skalic M, Elliott DJ, et al. SUPPA2: fast, accurate, and uncertainty-aware differential splicing analysis across multiple conditions. Genome Biol. 2018;19: 40.
